# REPARATION: Ribosome Profiling Assisted (Re-)Annotation of Bacterial genomes

**DOI:** 10.1101/113530

**Authors:** Elvis Ndah, Veronique Jonckheere, Adam Giess, Eivind Valen, Gerben Menschaert, Petra Van Damme

**Affiliations:** VIB-UGent Center for Medical Biotechnology, B-9000 Ghent, Belgium; Department of Biochemistry, Ghent University, B-9000 Ghent, Belgium; Lab of Bioinformatics and Computational Genomics, Department of Mathematical Modelling, Statistics and Bioinformatics, Faculty of Bioscience Engineering, Ghent University, B-9000 Ghent, Belgium; Computational Biology Unit, Department of Informatics, University of Bergen, Bergen 5020, Norway; Sars International Centre for Marine Molecular Biology, University of Bergen, 5008 Bergen, Norway

**Keywords:** ribosome profiling, machine learning, N-terminomics, bacteria, genome (re-)annotation, open reading frames

## Abstract

Prokaryotic genome annotation is highly dependent on automated methods, as manual curation cannot keep up with the exponential growth of sequenced genomes. Current automated methods depend heavily on sequence context and often underestimate the complexity of the proteome. We developed REPARATION (RibosomeE Profiling Assisted (Re-)AnnotaTION), a *de novo* algorithm that takes advantage of experimental protein translation evidence from ribosome profiling (Ribo-seq) to delineate translated open reading frames (ORFs) in bacteria, independent of genome annotation. REPARATION evaluates all possible ORFs in the genome and estimates minimum thresholds based on a growth curve model to screen for spurious ORFs. We applied REPARATION to three annotated bacterial species to obtain a more comprehensive mapping of their translation landscape in support of experimental data. In all cases, we identified hundreds of novel (small) ORFs including variants of previously annotated ORFs. Our predictions were supported by matching mass spectrometry (MS) proteomics data, sequence composition and conservation analysis. REPARATION is unique in that it makes use of experimental translation evidence to perform *de novo* ORF delineation in bacterial genomes irrespective of the sequence context of the reading frame.

## INTRODUCTION

In recent years, the advent of next generation sequencing has led to an exponential growth of sequenced prokaryotic genomes. As curation based methods cannot keep pace with the increase in the number of available bacterial genomes, researchers have reverted to the use of computational methods for prokaryotic genome annotation (Richardson and Watson 2013; Land et al. 2015). However, advanced genome annotation should entail more than simply relying on automatic gene predictions or transferred genome annotation, as these often introduce and propagate inconsistencies (Richardson and Watson 2013). Moreover, the dependence on sequence contexts of the open reading frame (ORF) by automatic methods often introduces a bias in gene prediction, as studies have shown that translation can occur irrespective of the sequence composition of the ORF (Michel et al. 2012; Fields et al. 2015). Further, gene prediction methods that depend solely on the genomic template often lack the capabilities to capture the true complexity of the translation landscape (Fields et al. 2015), overall stressing the need for non *in silico* based gene prediction approaches.

Ribosome profiling (Ingolia et al. 2009) (Ribo-seq) has revolutionized the study of protein synthesis in a wide variety of prokaryotic and eukaryotic species. Ribo-seq provides a global measurement of translation *in vivo* by capturing translating ribosomes along an mRNA. More specifically, ribosome protected mRNA footprint (RPFs) are extracted and converted into a deep sequencing cDNA library. When aligned to a reference genome, these RPFs provide a genome-wide snapshot of the positions of translating ribosomes along the mRNA at the time of the experiment (Ingolia et al. 2009). This genome-wide positional information of translating ribosomes allows for the identification of translated regions.

With the advent of Ribo-seq, numerous computational methods have been developed to detect putatively translated regions in eukaryotes, all taking advantages of inherent Ribo-seq based metrics to identify translated ORFs. In the studies of Lee et al. (2012) and Crappé et al. (2014), a rule based peak detection algorithm was used to identify translation initiation sites, while Bazzini et al. (2014) and Calviello et al. (2015) take advantage of the triplet periodicity property of Ribo-seq data. Fields et al. (2015) and Chew et al. (2013) developed an ensemble classifier that aggregate multiple features to predict putative coding ORFs. In addition to these tools Michel et al. (2016) developed a quality control toolbox available from Galaxy (RiboGalaxy) and Chung et al. (2015) developed a method to infer frameshifting events within coding regions. However, all these methods focus mainly on eukaryote genomes with a pre-defined transcriptome and are not directly transferable to prokaryotes genomes. Thus far, no computational method has yet been reported to systematically delineate ORFs in prokaryotic genomes based on Ribo-seq data. In this work, we aimed at developing an algorithm that makes use of experimental evidence of translation from Ribo-seq to perform *de novo* ORF delineations in prokaryotic genomes.

Our algorithm, REPARATION (RibosomeE Profiling Assisted (Re-)AnnotaTION) trains an ensemble classifier to learn Ribo-seq patterns from a set of confident protein coding ORFs for a *de novo* delineation of ORFs in bacterial genomes. REPARATION deduces intrinsic characteristics from the data and thus can be applied to Ribo-seq experiments targeting elongating ribosomes. We evaluated the performance of REPARATION on three annotated bacterial species. REPARATION was able to identify a multitude of putative coding ORFs corresponding to previously annotated protein coding and non-protein coding regions, variants of annotated ORFs (i.e. in-frame truncations or 5′ extensions) and intergenic ORFs. Further, we validated our findings using matching proteomics, sequence composition and phylogenetic conservation analyses and compared to other methods in the field.

## MATERIAL AND METHODS

REPARATION performs *de novo* ORF delineation by training a random forest classifier to learn Ribo-seq patterns exhibited by protein coding ORF. A random forest model was chosen over other algorithms for training because of its robustness to outliers, low bias and optimal performance with few parameter tuning (Hastie and Tibshirani 2009). The REPARATION pipeline (*figure 1A*) starts by traversing the entire prokaryotic genome sequence to generate all possible ORFs that have an arbitrary length of at least 10 codons (30nt) and initiate with either an ATG, GTG or TTG codon (the most frequently used start codons in a variety of prokaryotic species (Panicker, Browning, and Markham 2015)) until the next in-frame stop codon.

**Figure 1:**
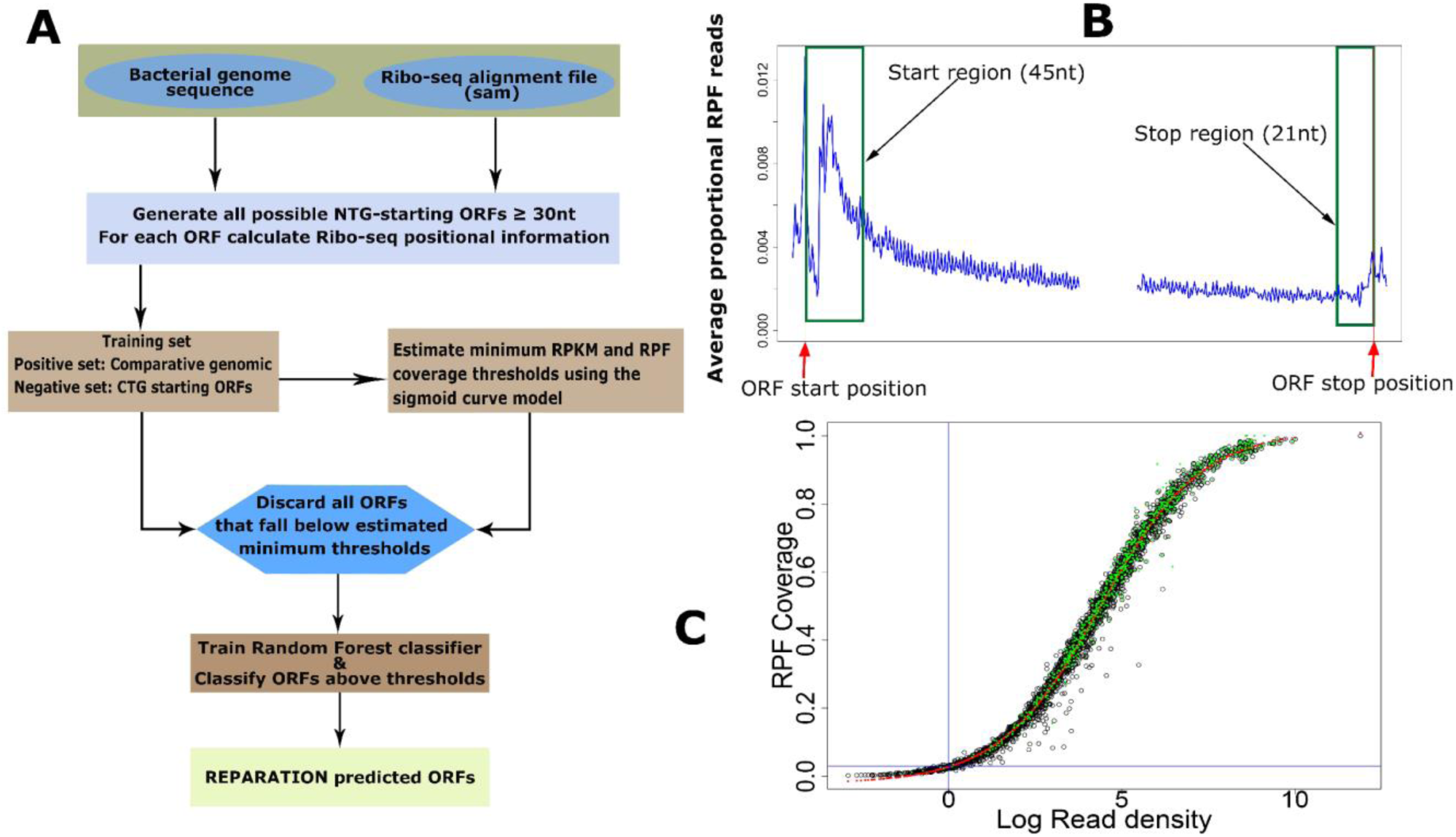
REPARATION pipeline for de novo ORF delineation in prokaryotes. *(A) REPARATION workflow diagram. The entire prokaryotic genome is traversed and all possible NTG-starting ORFs are generated. Next, ORF-specific positional Ribo-seq signal information is calculated based on the metagenic gene profile (B). To discard spurious ORFs, the minimum log2 RPKM and ORF RPF coverage thresholds are estimated using a four-parameter logistic S-curve (C). (B) Metagenic profile of salmonella data indicating read accumulation at the start and stop of ORFs (stitched together in the middle for visualization purposes). (C) S-curve with fitted four parameters logistic curve (red) and indication of predicted ORFs with support from N-terminal proteomics data (green) in the case of E. coli.*

### Training Sets

The set of positive examples is constructed by a comparative genomic approach. The algorithm uses Prodigal V2.6.3 (Hyatt et al. 2010) to generate an ORF set, this set is then BLAST searched against a database of curated bacterial protein sequences (e.g. UniprotKB-SwissProt). The BLAST search is performed using the UBLAST algorithm from the USEARCH package (Edgar 2010). ORFs that match at least one known protein coding sequence with a minimum e-value of 10^-5^ and a minimum identity of 75% are selected for the positive set. The negative set consist of ORFs starting with CTG viewing their infrequent occurrence as translation starts (<0.01%) in the annotations of the interrogated species (*supplementary table T1)* and at least the length of the shortest ORF in the positive set. We then grouped all CTG ORFs sharing the same in frame stop codon into an “ORF family”. Per ORF family we select the longest ORF as a representative member of that “ORF family”.

### Feature construction

To train the random forest classifier we constructed five features based on Ribo-seq signals of translated ORFs and complemented these with the ribosome binding energy measurements (Suzek et al. 2001). The meta gene profile shown in *figure 1B* illustrates a Ribo-seq signal pattern reminiscent to patterns previously reported for protein coding transcripts in prokaryotes for Ribo-seq experiments that targets elongating ribosomes (Woolstenhulme et al. 2015). The profile exhibits read accumulation within the first 40-50nts downstream of the start and a slight increase just before the stop codon. The features used in the model are as follows:

1. Start and stop region read density (RPKM). We defined a start region of an ORF by taking 3nt upstream (to account for any error in P-site assignment) and 45nt downstream of the ORF start position, while the stop region constitutes the last 21nt upstream of the stop position. Of note, for ORFs shorter than 63 nucleotides we used the first 70% and last 25% of the ORF length to model the start and stop regions of the ORF. The ORF RPF read count per nucleotide position is divided by the total RPF reads within the ORF to ensure that features are comparable across different ORFs. The start and stop region RPF read densities are subsequently calculated from the proportional reads.
2. ORF coverage and start RPF coverage. We defined the ORF (start) RPF coverage as the proportion of nucleotide positions covered by RPF reads within the entire ORF (start region). RPF coverage is calculated from the RPF positional read profile.
3. Read accumulation proportion. This feature measures the ratio of the RPF reads accumulated at the start region (first 45nt) relative to RPF reads within the rest of the ORF. It is defined by equation 1

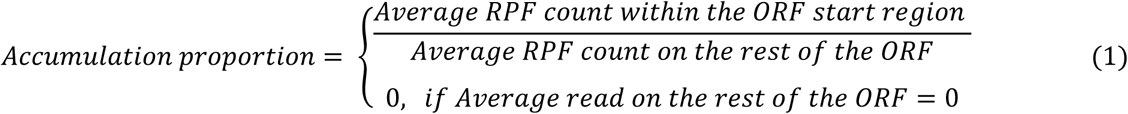 We reasoned that since Ribo-seq reads tend to accumulate within the start region of a translated ORF relative to the rest of the ORF, correctly delineated ORFs will tend to have score greater than one. Spurious ORFs that overlap at the start or stop of translated ORFs will score lower as their non-overlapping regions would tend to have no reads, hence resulting to accumulation scores less than one.
4. Ribosome binding site (RBS) energy. The interaction between Shine-Dalgarno (SD) sequence and its complementary sequence in the 16S rRNA (anti-SD), referred to as SD ribosome binding site (RBS) was proven to be very important in the recruitment of the ribosome for translation initiation in bacteria (Shultzaberger et al. 2001). As such, and to aid in the prediction of SD/anti-SD dependent translation events, the ribosome’s free binding energy or ribosome binding site (RBS) energy was included as in feature in the model. The RBS energy, representative of the probability that the ribosome will bind to a specific mRNA and thus proportional to the mRNA’s translation initiation rate, was calculated using the distance dependent probabilistic method and using the anti-Shine Dalgarno (aSD) sequence GGAGG as described in Suzek et al. (2001).The inclusion of the of the RBS energy features in the prediction model as well as the aSD sequence are user defined parameters to allow for bacterial species where non aSD/SD dependent translation events have been reported (Shultzaberger et al. 2001; Hyatt et al. 2010; Omotajo et al. 2015).

### Sigmoid (S)-curve model

Since REPARATION pipeline was developed to allow for ORFs as short as 30nt, this results in an exponential increase of potential ORFs. To ensure the algorithm is traceable, we defined minimum threshold values to eliminate spurious ORFs. To do this we take advantage of the sigmoid curve (S-curve) relationship observed between ORF RPF coverage and the ORF log2 read density (RPKM) as depicted in *figure 1C and supplementary figure S1*. The fitted logistic curve (red), modelled by a 4-parameter logistic regression (*equation 2*) and describing the relationship between ribosome density and RPF coverage was used to estimate the minimum read density and ORF RPF coverage to allow for correct ORF delineation. We estimated the lower bend point of the fitted 4 parameter logistic regression using the method described in (Lutz and Lutz 2009) and implemented in the R Package *Sizer* (Sonderegger et al. 2009).

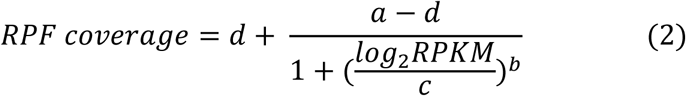

Where *d* represents the ORF coverage for infinite read density (RPKM) and *a* the minimum ORF coverage at RPKM zero while *b* and *c* represents the slope of the curve and the RPKM value at *(d – a)/2* respectively.

### Post Processing Random Forest predicted ORFs

We implement a rule based post processing algorithm to eliminate false positives that might be called because they share overlapping regions with actual coding ORFs (*supplementary figure S2*). First, considering the simplified assumption that bacterial genes can have only one possible translation start site, we group all predicted ORFs sharing the same in frame stop codon into an “ORF family”. *Supplementary figure S2 A* depicts an ORF family with two predicted starts, if start S1 has more reads than S2 then we select S1 as the gene start. If there are no Ribo-seq reads between S1 and S2 then we select S2 as the gene start since S1 adds no extra information to the gene profile. If S1 has more reads than S2 but if S1 falls within the coding region of an out-of-frame upstream predicted ORF on the same strand, we select S1 as the most likely start if there is a peak (i.e. kurtosis > 0) within a window of -21 to +21 around S1.

Next we consider two overlapping ORFs on different frames as depicted in *supplementary figure S2 B*. If the read density and RPF coverage of the non-overlapping region of F1 are less than the S-curve estimated thresholds, then F1 is dropped in favor of F2 and vice versa. If both non-overlapping regions have a read density and RPF coverage greater than the minimum, then we assume both are expressed. Finally, we drop all internal out-of-frame ORFs falling completely within another ORF (supplementary *figure S2 C).*

## RESULTS

To assess the performance and utility of our REPARATION algorithm (figure 1A), besides two publicly available bacterial Ribo-seq datasets from Escherichia coli K12 str. MG1655 and Bacillus subtilis subsp. subtilis str. 168 (Li et al. 2014), we generated ribosome profiling data and matching RNA-seq data from a monosome and polysome enriched fraction (Heyer & Moore 2016) of Salmonella enterica serovar Typhimurium strain SL1344 (experimental details in supplementary methods).

REPARATION starts by traversing the entire prokaryotic genome sequence to generate all possible ORFs that have an arbitrary length of at least 10 codons (30nt) and initiate with either an ATG, GTG or TTG codon (the most frequently used start codons in a variety of prokaryotic species (Panicker et al. 2015)) until the next in-frame stop codon. REPARATION applies a random forest classifier trained on features derived from the meta-gene profile of known protein coding ORFs (figure 1B). These features encompass 1) the start region (first 45nt of the ORF) read density, 2) the stop region (last 21nt) read density (Fields et al. 2015) 3) ORF RPF coverage refers to the proportion of nucleotides within the ORF covered by positional RPF reads (Chew et al. 2013), 4) start region RPF coverage, i.e. the proportion of nucleotides within the start region covered by RPF reads, 5) the ratio of the average RPF read count within the start region divided by the average RPF read count within the rest of the ORF and 6) ribosome binding site (RBS) energy (see material and methods). The classifier’s training set consisted of positive examples generated using a comparative genomic approach. First we used prodigal (Hyatt et al. 2010) to generate a set of ORFs, which were subsequently BLAST searched against a curated set of bacterial protein sequences from UniProtKB-SwissProt, ORFs with e-values less than 10^-5^ and a minimum identity score of 75% were retained. Viewing their infrequent occurrence as translation starts (<0.01%) in the annotations of the interrogated species, the negative set consisted of all CTG-starting ORFs (*supplementary table T1*). The algorithm then estimates a minimum read density and ORF RPF coverage to discard spurious ORFs by exploiting the sigmoid relationship between these features (figure 1C). Using a four-parameter logistic regression curve on the positive set, REPARATION estimates the lower bend point of the fitted curve representing a two-dimensional threshold (read density and ORF RPF coverage) (*supplementary table T2*). All ORFs with read density and ORF RPF coverage below these thresholds were discarded, including those in the training set. When trained on these sets, the random forest classifier achieved on average an 89, 90 and 92% 10-fold cross validation accuracy with area under the curve values of 0.92, 0.92 and 0.93 for the Salmonella, E. coli and Bacillus data sets respectively (*supplementary figure S3 A*).

Of the three species evaluated, REPARATION mapped putative coding ORFs corresponding to regions annotated as protein coding, as well as to non-coding and intergenic.

### REPARATION-predicted ORFs predominantly match to, or overlap with annotated ORFs and follow the reference model of start codon usage

Viewing the previously reported similarities in the translation properties of monosomes and polysomes (Heyer & Moore (2016)) and the high correlation observed between the two samples (*supplementary figure S3 B*), we considered the *Salmonella* monosome and polysome samples as replicate samples for the purpose of translated ORF delineation.

For *Salmonella*, REPARATION predicted a total of 3957 and 3881 putative ORFs in the monosome and polysome sample respectively. Of these, 3421 (88%) ORFs found common in both datasets were considered as the high confident ORF set (*supplementary file F1*). For *E. coli*, a high confident set of 3202 (90%) was selected based on the 3594 and 3569 predicted ORFs in replicate samples 1 and 2 respectively (*supplementary file F2*). Thirdly, in the *Bacillus* sample, 3435 putative coding ORFs were predicted (*supplementary file F3*).

From the high confident set of predicted ORFs in *Salmonella* and *E. coli*, 82% (2822) and 89% (2855) correspond to previously annotated ORFs (respectively), while 80% (2733) of the *Bacillus* predicted ORFs corresponds to previously annotated ORFs (*figure 2*). 14, 8 and 14% of predicted ORFs in the *Salmonella, E. coli and Bacillus* samples (respectively), correspond to variants of previously annotated ORFs, potentially giving rise to N-terminally truncated or extended protein variants referred to as N-terminal proteoforms (Gawron, Gevaert, and Damme 2014). Consequently, in *Salmonella* and *E. coli* 3% belong to novel putative coding regions while in *Bacillus* 7% belong to novel ORFs.

**Figure 2:**
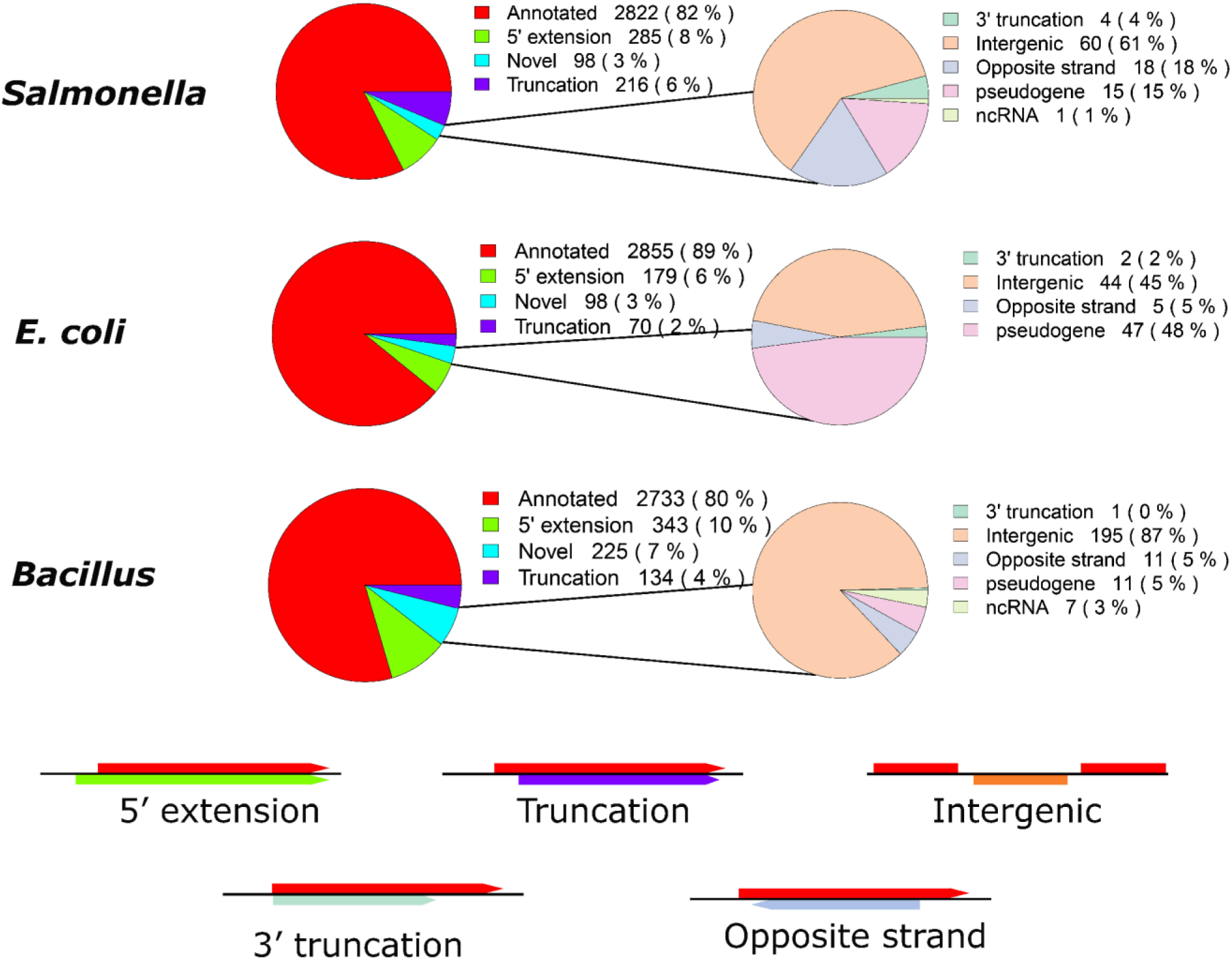
*Proportion of REPARATION predicted ORFs per ORF category for the high confident ORF sets in Salmonella and E. coli as well as for the Bacillus predictions.* Overall, 71%, 74% and 77% (including the variants) of all *ENSEMBL* annotated protein coding ORFs in *Salmonella, E. coli* and Bacillus (respectively) were predicted by REPARATION.

On average the truncations were 26, 26 and 51 codons downstream of the annotated starts while the extensions where 18, 13 and 9 codons upstream for *Salmonella, E. coli* and *Bacillus* (respectively). Of note, 60, 53 and 77 of the predicted variants display only a 1 codon shift from the annotated starts in *Salmonella, E. coli* and *Bacillus* respectively (*supplementary table T3*).

In our evaluation of REPARATION, we allow for the three commonly used start codons in prokaryotes ATG, GTG and TTG as translation initiation triplet. Of note however, REPARATION was designed without any bias in start codon selection for ORF prediction. The hierarchy of start codon usage over all predicted ORFs are consistent with the standard model for translation initiation in the *ENSEMBL* annotation of the corresponding species i.e. in case of *Salmonella* and *E. coli*, a preference of ATG over GTG and TTG and ATG>TTG>GTG in *Bacillus* (*table 1*) could be observed.

**Table 1:** Start codon usage distribution of the predicted putative coding ORFs. *The predicted ORFs in all three species follow the starts codon usage distributions of the corresponding species annotation. In case of Salmonella* and *E. coli, only ORFs from the high confident set were considered.*

|  | **ENSEMBL**<br>**Annotation** | **All**<br>**predictions** | **Matching**<br>**Annotated** | **Extensions** | **Truncations** | **Novel** |
| --- | --- | --- | --- | --- | --- | --- |
| ***Salmonella*** |  |  |  |  |  |  |
| ATG | 4093 (88.0%) | 2942 (86%) | 2576 (91.2%) | 166 (58%) | 139(64%) | 61 (62%) |
| GTG | 429 (9.20%) | 344 (10%) | 206 (7.4%) | 64 (24%) | 52(24%) | 18 (18%) |
| TTG | 126 (2.70%) | 135 (4%) | 40 (1.4%) | 51 (18%) | 25(12%) | 19 (20%) |
| ***E. coli*** |  |  |  |  |  |  |
| ATG | 3747 (90.1%) | 2776 (87%) | 2591 (91%) | 76 (42%) | 49 (70%) | 60 (62%) |
| GTG | 386 (9.2%) | 284 (9%) | 209 (7%) | 44 (25%) | 9 (13%) | 22 (22%) |
| TTG | 71 (2.0%) | 142 (4%) | 55 (2%) | 59 (33%) | 12 (17%) | 16 (16%) |
| ***Bacillus*** |  |  |  |  |  |  |
| ATG | 3253(77.7%) | 2502 (73%) | 2176 (80%) | 141 (41%) | 75 (56%) | 110(49%) |
| GTG | 386 (9.2%) | 413 (12%) | 237 (9%) | 108 (31%) | 29 (22%) | 39 (17%) |
| TTG | 529 (12.6%) | 520 (15%) | 320 (12%) | 94 (29%) | 30 (22%) | 76(34%) |

Interestingly however, we observe that novel and variant ORFs are enriched for being initiated at near-cognate start codons when compared to annotated ORFs. In case of the variants, this bias is most likely due to the preference of automatic gene prediction methods to select a neighbouring ATG as the start codon (Salzberg et al. 1998; Hyatt et al. 2010).

### Novel ORFs are evolutionary conserved and display similar amino acid sequence patterns as compared to annotated ORFs

To gain insight into the novel predictions, we analysed and compared their evolutionary conservation pattern to that of predicted annotations. Novel and extended ORFs exhibit similar conservation patterns to annotated ORFs, with higher nucleotide conservation from the start codon onwards and within the upstream ribosomal binding site or Shine Dalgarno region positioned -15 to -5nt upstream of the predicted start (*figure 3*), a region aiding in translation initiation by its base pairing with the 3’-end of rRNA (Shultzaberger et al. 2001; Suzek et al. 2001). The higher conservation and triplet periodicity observed upstream of the truncations is likely because in some cases multiple forms of the gene (i.e. N-terminal proteoforms) are (co-)expressed (*supplementary table T4*). A manual inspection of the alignments indeed indicates that different forms of the genes are expressed across different species. Of the 66 truncations used in the *Salmonella* conservation analysis, 45% shows evidence of the existence of multiple forms across different bacterial species, while in case of *E. coli* and *Bacillus* these percentages were 40 and 28 from 26 and 25 truncations respectively.

**Figure 3:**
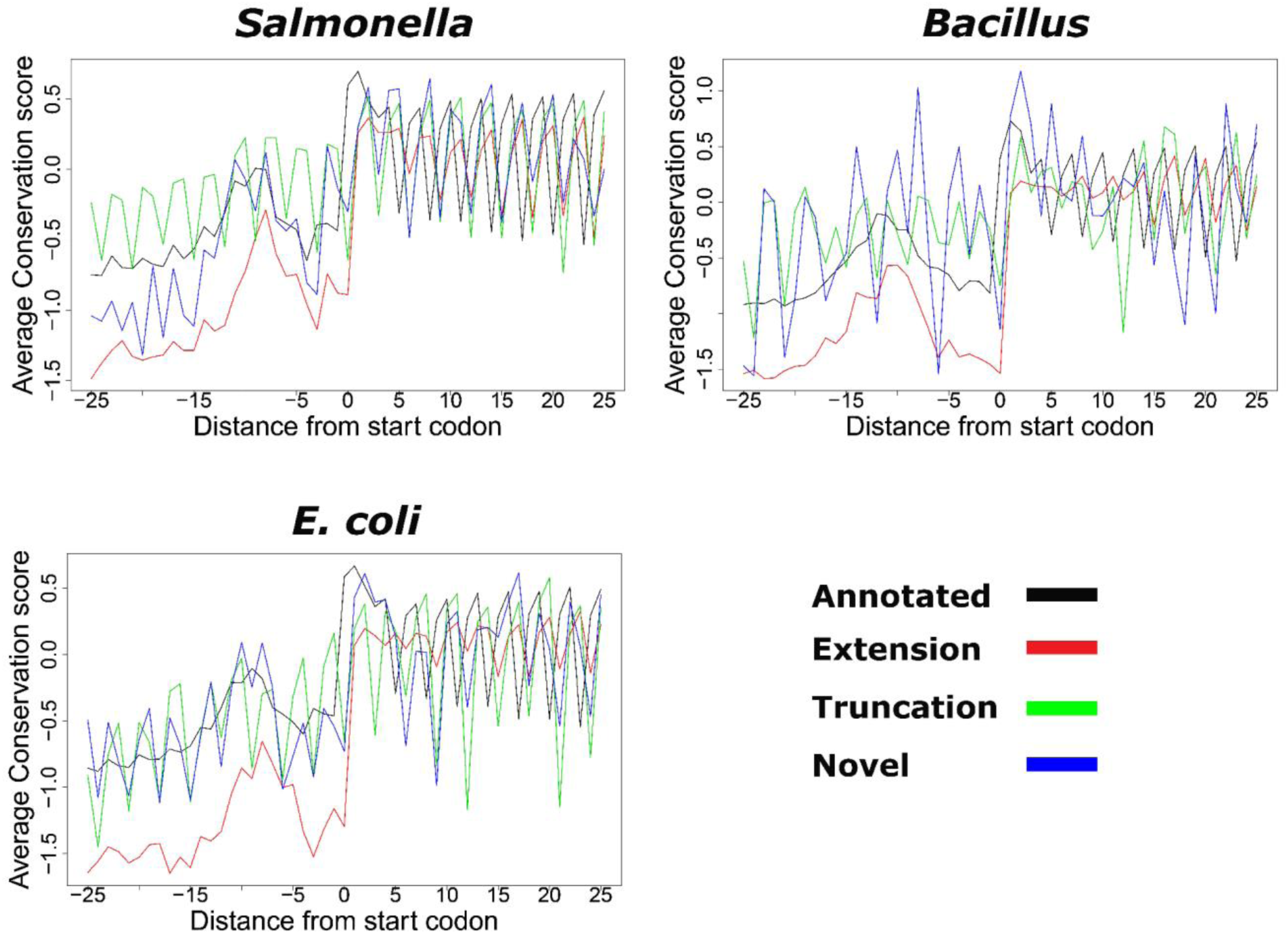
*Conservation pattern of REPARATION predicted ORFs. Nucleotide conservation scores are calculated using the Jukes cantor conservation matrix for nucleotides. Site conservation scores are calculated using the rate4site algorithm and displayed for a +/- 25nt window around the predicted start site. The site conservation score was calculated only for ORFs with at least 5 orthologous sequences from a collection of randomly selected bacteria protein sequences from species within the same family as Salmonella/E. coli and Bacillus and outside the family. 833 annotated, 161 extensions, 99 truncations and 12 novel ORFs had at least 5 orthologous sequences in case of Salmonella, while the E. coli profile consisted of 2359 annotated ORFs, 70 extensions, 26 truncations and 18 novel ORFs were considered. In the case of Bacillus there are 1886 annotated, 112 extensions, 19 truncations and 2 novel ORFs*

Of the 98 novel ORFs predicted in *Salmonella*, 48% (47) had at least one reported orthologous sequence (*supplementary file F1*). While 59% (58 out of 98) and 19% (42 out of 225) in *E. coli* and *Bacillus* (respectively) had at least one orthologous sequence (*supplementary files F2 & F3*).

To further confirm that the newly identified ORFs do not represent random noise, we compared the amino acid composition of predicted annotations to that of novel putative coding ORFs and to a set of randomly generated amino acid sequences of equal lengths to predicted ORFs matching annotations. In all three species, we observe a very high correlation (≥0.80) between the amino acid compositions of novel and annotated ORFs (*supplementary figure S4*). While a generally poor correlation (≤0.19) was observed when comparing novel or annotated ORFs against a random set of ORFs.

Since evolutionarily conserved significant biases in protein N- and C-termini were previously reported for pro- as well as eukaryotes, often with pronounced biases at the second amino acid positions (van Damme et al. 2011; Palenchar 2008), we next investigated whether the amino acid usage frequency at second position of the novel and re-annotated ORFs exhibited a similar pattern to that of annotated ORFs. Compared to amino acid frequency in the species matching proteomes, clearly the overall distribution is similar for the two ORF categories. More specifically, a significant enrichment of Lys (about 3-fold) at the second amino acid position was observed in case of all three species analysed. For *Salmonella* and *E. coli*, Ser and Thr at the second amino acid position was equally enriched while in case of *Bacillus*, Asn was slightly more frequent in the second position while other amino acids are clearly underrepresented (i.e. Trp and Tyr) (*supplementary figure S5*), observations very well in line with previous N-terminal biases observed (Palenchar 2008).

### Proteomics assisted validation of REPARATION predicted ORFs

To validate our predicted ORFs we generated N-terminal and shotgun proteomics data from matching *E. coli* and *Salmonella* samples respectively. While N-terminomics enables the isolation of N-terminal peptides, making it appropriate for the validation of translation initiation events, shotgun proteomics provides a more global assessment of the expressed proteome. Three different proteome digestions were performed in the shotgun experiment to increase proteome coverage. The shotgun and N-terminal proteomics data were searched against a six-frame translation database of the *E. coli* and *Salmonella* genomes. In both experiments, and based on identified peptides, longest non-redundant peptide sequences were aggregated to map onto the REPARATION predictions.

In case of Salmonella, 10751 unique peptides belonging to 2235 ORFs in the six-frame translation database were identified by means of shotgun proteomics. Of these, 92% (9891) correspond to 1794 REPARATION predicted ORFs (*figure 4A*), the 9% missed by REPARATION mostly represent lowly expressed ORFs (*Supplementary figure S6 A*). While most shotgun peptides support previously annotated regions (*figure 4B*), we additionally identified peptides in support of novel ORFs and ORF reannotations (i.e. N-terminal protein extensions). More specifically, supportive evidence was found in the case of 8 novel ORFs and 21 extensions having at least one identified peptide with a start position upstream of the annotated start (*supplementary file F1*).

**Figure 4:**
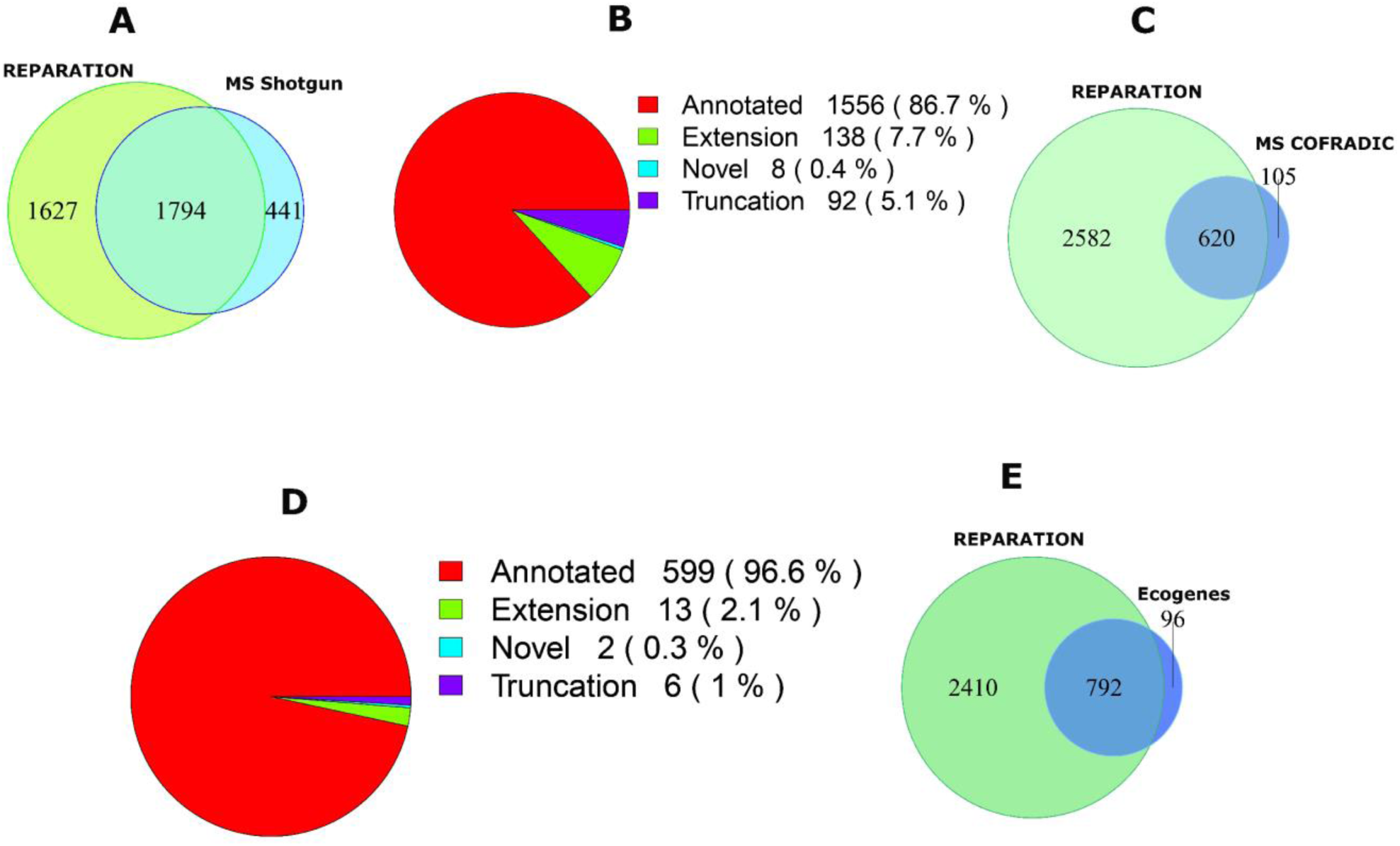
*MS validation of REPARATION pipeline. A) Overlap between the protein sequences identified from shotgun proteomics and the REPARATION predicted ORFs in Salmonella. B) The number of ORFs per category with at least one identified peptide for the high confident set of Salmonella predicted ORFs. C) Overlap between ORFs with N-terminal peptide support and REPARATION predicted ORFs in E. coli. D) Number of predicted ORFs for each category with N-terminal peptide support in the E. coli high confident set. E) Overlap between REPARATION predicted ORFs and the Ecogene verified E. coli ORFs.*

For *E. coli,* N-terminal proteomics identified a total of 785 blocked N-terminal peptides that are compliant with the rules of initiator methionine processing (see *supplementary methods*) belonging to 781 ORFs. Under the assumption that none of these ORFs have multiple initiation sites we choose the most upstream N-terminal peptide and overlapped these with the REPARATION predictions. Of the 781 ORFs with peptide support 725 pass the S-curve estimated minimum thresholds. 86% (620) of these matched REPARATION predicted N-termini (*figure 4C & D*), while in 6% of the cases, a different translation start was predicted by REPARATION downstream in 10 cases (with an average distance of 8 codons) and upstream in 40 cases (with an average distance of 39 codons) from the TIS matching the identified N-terminal peptides. The remaining 8%, not predicted by REPARATION, mainly represent N-termini originating from lowly expressed ORFs (*supplementary figure S6 B*). Most of the N-terminal supported ORFs matched annotations, while 21 correspond to re-annotations or novel ORFs (13 extensions, 6 truncations and 2 novel). We also assessed the predicted ORFs against the 917 *E. coli K-12 Ecogenes* verified protein coding sequences, a set consisting of proteins sequences with their mature N-terminal residues sequenced using N-terminal Edman sequencing (Krug et al. 2013). Of these, 893 pass the estimated minimum thresholds of which 89% (792) matched REPARATION predicted ORFs (*figure 4E*). REPARATION predicted a different start sites in case of 54 of *Ecogene* verified ORFs, including 45 upstream (average distance of 8 codons) and 9 downstream TISs (average distance of 35 codons).

### REPARATION in the aid of genome re-annotation

In case of the three species-specific translatomes analysed, REPARATION uncovered novel putative coding genes in addition to extensions and truncations of previously annotated genes with supporting proteomics and conservation evidence. More specifically, in the case of the gene *adhP (Salmonella*), REPARATION predicts that translation initiates 27 codons upstream of the annotated start, an ORF extension supported by an N-terminal peptide identification (*figure 5A*) and of which the corresponding sequence is conserved (*supplementary figure S7 A*). N-terminal peptide support, next to the clear lack of Ribo-seq reads in the region between the novel and annotated start (*figure 5B)* of gene *yidR* (*E. coli*), also points to translation initiating 11 codons downstream of the annotation start as predicted by REPARATION. A novel putative coding gene was found matching the intergenic region *Chromosome:2819729-2820319* (*Salmonella)* with Ribo-seq and RNA-seq signals complemented by two unique peptide identifications (*figure 5C*).

**Figure 5:**
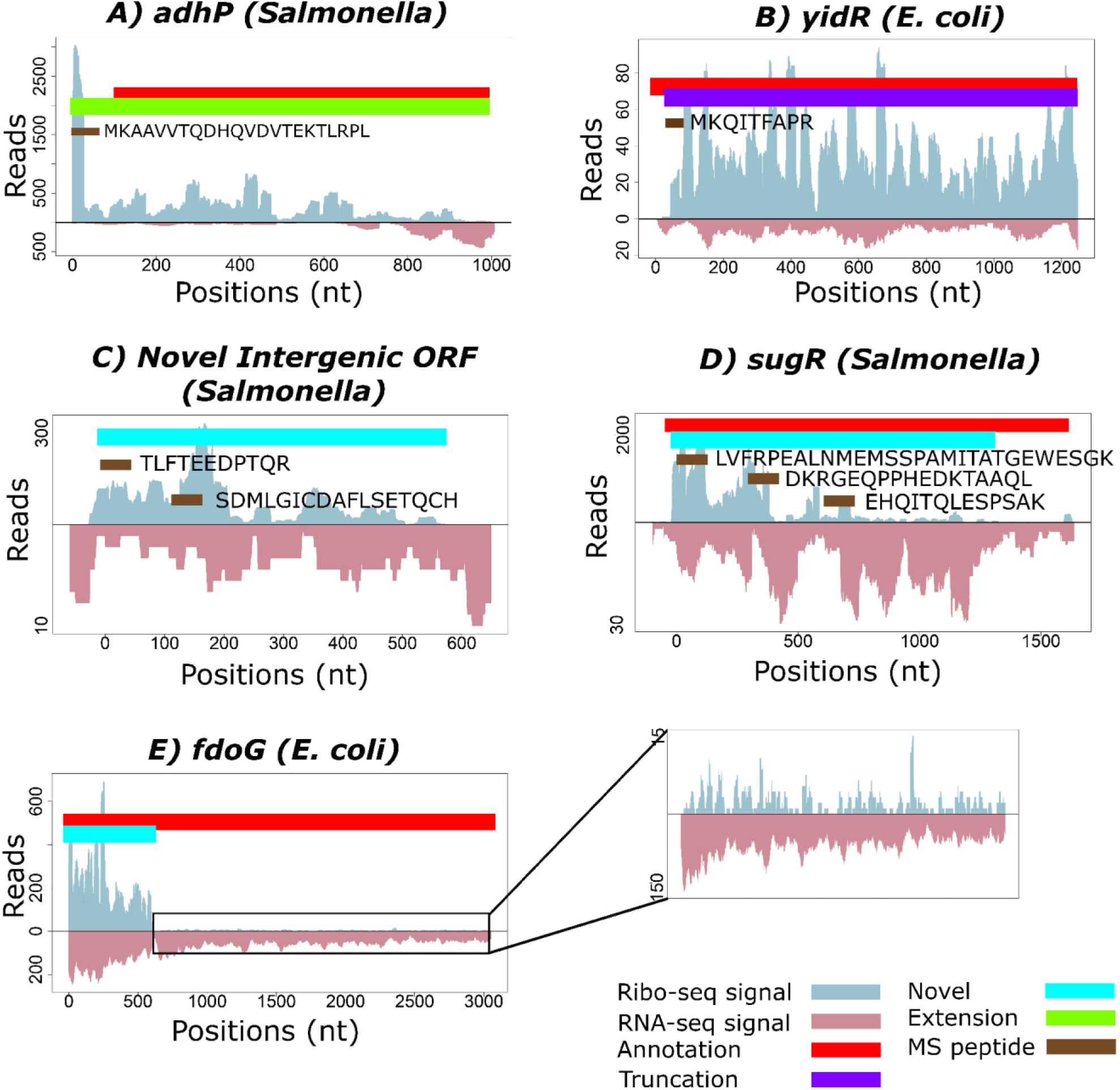
REPARATION assisted reannotation of bacterial genomes. *A) REPARATION predicted 5′ extension of the adhP gene (Salmonella) with supportive peptide evidence mapping upstream and in-frame with the annotated ORF. B) Gene yidR (E. coli) predicted as a 5′ truncation with N-terminal peptide support and Ribo-seq reads starting downstream of the annotated start. C) Novel putative coding intergenic ORF in the region Chromosome:2819729-2820319 (Salmonella) with supportive peptide evidence. D) Evidence of translation within pseudogene sugR (Salmonella), with two matching peptide identification. E) Putative co-expression of a 3′ truncated ORF as well as its 3′ extended counterpart due to a frameshifting event occurring during translation of the fdoG gene (E. coli). A magnification of the region beyond the stop codon displays a continuous, though ∼100-fold lower pattern of Ribo-seq reads indicative of stop codon read-through.*

Of note, there are currently 72, 182 and 70 annotated pseudogenes in the current *ENSEMBL* annotations of *Salmonella, E. coli* and *Bacillus* (respectively). REPARATION predicted conserved putative coding ORFs within 12, 35 and 11 pseudogene regions leading to 15, 47 and 11 predicted ORFs for *Salmonella, E. coli* and *Bacillus* (respectively). Since pseudogenes in bacteria are typically modified/removed rapidly during evolution, coupled with the fact that only uniquely mapped reads were allowed, the observed conservation with the existence of functional orthologues points to the genuine coding potential of these loci and thus functional importance of their translation product (Goodhead and Darby 2015; Lerat and Ochman 2005). One representative example is the identified putative coding ORF in the *sugR* pseudogene (*Salmonella*) which is supported by 3 unique peptide identifications (*figure 5D*).

Interestingly, in case of *fdoG (E. coli)*, REPARATION predicted two juxta positioned ORFs, both contained within a previously annotated ORF harbouring a stop codon read through event (*figure 5E*). In *E. coli* three other such read through events have been reported for genes *fdnG*, *fdhF* and *prfB*. The Ribo-seq read density within the C-terminal truncated ORF is about 100-fold higher as compared to the 5′ truncated ORF while only a 3-fold difference in RNA-seq density could be observed. The RNA-seq evidence supports the presence of the stop codon TGA at the end of the C-terminal truncated ORF. However, the observed continuous Ribo-seq signal is indicative of translation beyond the sequencing-verified stop codon and most likely points to a stop codon read through event (Feng et al. 2012). The so-called 3′ and 5′ truncations of the current annotation predicted by REPARATION are likely due to the algorithm not allowing for stop codon read through. A similar trend was observed in case of its *Salmonella* orthologue, with the Ribo-seq signal and RNA-seq 30- and 2-folds (respectively) higher for the C-terminal truncated ORF than the longer N-terminal truncated ORF (*supplementary figure S7 B*).

### REPARATION in the aid of small ORF annotation

Small ORFs have historically been ignored in most *in silico* predictions because of the assumption that they can easily occur by chance due to their small size (Hyatt et al. 2010). As 71 codons is the average length when considering the length of the 5% shortest annotated ORFs in the 3 species, we here arbitrarily define a sORF as a translation product with a length of ≤71 codons. In *Salmonella*, REPARATION predicted 119 putative coding sORFs. Of these, 61 (51%) matched annotations and respectively 17 (14%) and 41 (34%) represent re-annotations (i.e. 3 extensions and 14 truncations) and novel ORFs. Supportive proteomics data was found for 29 predicted sORFs. While in *E. coli* and *Bacillus the* algorithm predicted 125 (95 (76%) matching annotations, 1 extension, 4 truncations and 26 novel) and 395 (161 (41%) matching annotations, 8 extensions, 29 truncations and 197 (50%) novel) sORFs. An interesting example of a possible re-annotation of gene *yfaD* (*E. coli*) is the REPARATION predicted 56 codon sORF representing a truncated form (*figure 6A*). In line with transcriptional data pointing to transcription of an mRNA not encompassing the annotated ORF, Ribo-seq indicates expression of a smaller ORF of which the start of the gene is located 243 codons downstream of the annotated start. Other representative examples are the intergenic 47 codons long sORF *Chromosome:2470500-2470643 (E. coli) (figure 6B),* the *30* codons long sORF located on the reverse strand of the *fre* gene (Salmonella) (*figure 6C*) and a 57 codon long intergenic *Bacillus* sORF that overlaps with the CDS of the sORF-encoding *hfq* gene (*figure 6D*).

**Figure 6:**
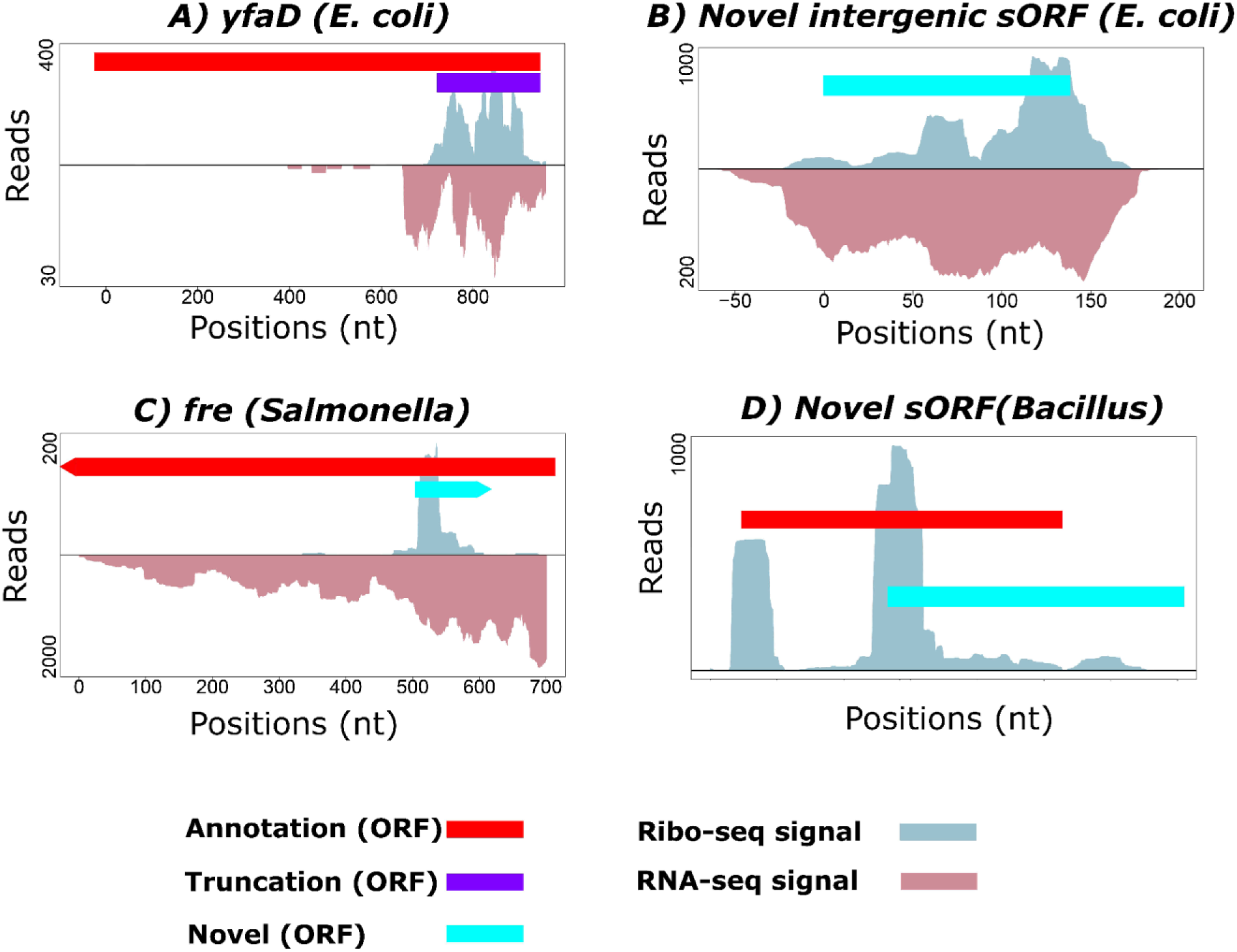
Novel sORFs predicted by REPARATION. *Ribo-seq and RNA-seq profiles indicate expression of A) a truncated form of the annotated yfaD (E. coli) gene B) a 47 codons sORF matching the region Chromosome:2470500-2470643 (E. coli.) C) a sORF encoded on the reverse strand encoding the fre gene (Salmonella). D) a sORF Chromosome:1867485-1867655 (Bacillus) that partially overlaps the annotated fhq sORF (Bacillus). The Ribo-seq profiles indicate translation initiation in another frame.*

## DISCUSSION

Experimental signals from ribosome profiling exhibit patterns across protein coding ORFs which can be exploited to accurately delineate translated ORFs. Although Ribo-seq is not completely standardized (Diament and Tuller 2016) and certain experimental procedures such as treatments (e.g. no treatment versus antibiotic treatment) tend to have a noticeable influence on the translation patterns observed (Calviello et al. 2015), we here developed an algorithm that enables a *de novo* delineation of translated ORFs in bacterial genomes. Our algorithm delineates putative protein coding ORFs in bacterial genomes using experimental information deduced from Ribo-seq, aiming to minimize biases inherent to *in silico* prediction methods.

Several methods that use ribosome profiling data to delineate open reading frames in eukaryotes have been reported, but due to peculiarities in prokaryotic genomes such as high gene density, occurrence of multiple overlapping genes (Huvet and Stumpf 2014) and the requirement of methodological adaptations to perform ribosome profiling experiments in bacteria, these tools are not directly transferable to ORF predictions in prokaryotes. Among them, and as cautioned by the authors, ORF-RATER is currently not applicable for use in prokaryotes as it performs its analysis on individual genes but prokaryotic genomes may not be divisible into distinct genes (Fields et al. 2015). Another method proposed by Chew et al. (2013) define metrics calculated from RPF read discrepancies between the 5′ and/or 3′ untranslated regions and the relative coding region of the transcript, metrics which cannot be properly defined in bacterial genomes due to their polycistronic nature. Alternative available software packages enabling the delineation of eukaryotic ORFs such as riboseqR (Chung et al. 2015) and RiboTaper (Calviello et al. 2015) take advantage of triplet periodicity in Ribo-seq data to infer translated ORFs. The former is a tool for parsing and inferring reading frames and transcript specific behaviour of Ribo-seq data while the later explores the triplet periodicity across the three frames within an annotated coding region to infer all possible coding ORFs. Due to the breakdown of triplet periodicity because of read accumulation at the start of the ORFs (*figure 1B*) coupled with the fact that only ATG starting ORFs are considered, applying RiboTaper on the Salmonella bacteria Ribo-seq data predominantly resulted in the prediction of 5′ truncated ORFs (*supplementary table T5*) and thus, while clearly some triplet periodicity can be observed, the strict reliance of RiboTaper on this feature makes it evidently not suited for TIS prediction and ORF delineation in bacteria. Further, all the afore mentioned tools heavily rely on an existing genome annotation and transcriptome structure which is often not available in the case of prokaryotic genomes. The limitations of the current existing methods coupled with the peculiarities of prokaryotic genome structures stresses the need for a dedicated method to delineate open reading frames in prokaryotes.

We applied REPARATION on three annotated bacterial species to illustrate its wide-applicability and ability to predict putative coding regions. Multiple lines of evidence, including proteomics data, evolutionary conservation analysis and sequence composition suggest that the REPARATION-predicted ORFs represent *bona fide* translation events. As expected, most predicted ORFs agreed with previous annotations, but additionally we could detect a multitude of ORF updates next to novel translated ORFs mainly within intergenic and pseudogene regions. While we clearly observed a shift towards near-cognate versus cognate start selection for the novel predictions, we nonetheless observe that the order of start codon usage follows the standard model in the respective species. Perhaps unsurprisingly viewing the difficulty to predict short ORF using classical gene predictions, the novel ORFs predicted by REPARATION are predominantly shorter than those previously annotated. Our predictions also point to possible errors in the current start site annotation of some genes, resulting in the identification of N-terminal truncations and extensions. The predicted extensions exhibit a similar conservation pattern to annotated ORFs while a higher conservation and triplet periodicity upstream of the truncated predictions (*figure 3*) is likely due to the expression of multiple proteoforms across species. The identification of multiple TIS-indicative N-termini in our *E. coli* N-terminomics dataset point to the existence of multiple translation initiation sites in at least 10 genes (*supplementary table T4*), likely an underrepresentation due to the low steady-state levels of N-terminally formylated N-termini. The former observation is in line with the recently revealed and until then highly underestimated occurrence of alternative translation events in eukaryotes (Ingolia, Lareau, and Weissman 2011; Van Damme et al. 2014). It is noteworthy that we identified 11 genes with multiple TIS-evidence from N-terminomics data (*supplementary table T4 and figure S4 C*), in case of REPARATION however, only a single ORF is selected per ORF family, therefore representing a bias toward the discovery of multiple proteoforms.

A substantial portion of the novel ORFs, with at least one identified orthologous gene, overlaps with known pseudogene loci. By virtue of the fact that pseudogenes in bacteria tend to be (sub)genus-specific and are rarely shared even among closely related species (Goodhead and Darby 2015; Lerat and Ochman 2005), it is likely that (part of) these genes have retained their protein coding potential, a finding that is further corroborated by proteomics data. The relatively fewer peptide identifications corresponding to the translation products of novel ORFs may in part be due to the difficulty of identifying these by MS, mainly because of their predominantly shorter nature and thus likely lower number of identifiable peptides (Fields et al. 2015). An *in-silico* analysis of the identifiable tryptic peptide coverage shows that on average 85% of the annotated protein sequences are covered by identifiable tryptic peptides while on average only 69% of the novel ORFs are covered by identifiable tryptic peptides. Furthermore, bacterial translation products of sORFs have previously been shown to be more hydrophobic in nature and therefore extraction biases might also (in part) contribute to their underrepresentation in our proteomics datasets (Hemm et al. 2008).

Historically sORFs have been neglected both in eukaryotes as well as prokaryotes. However, recently renewed interest has been directed toward the identification and characterization of sORFs (Andrews and Rothnagel 2014; Bazzini et al. 2014; Olexiouk et al. 2016). Small proteins represent a particularly difficult problem because they often yield weak statistics when performing computational analysis, making it difficult to discriminate protein coding from non-protein coding small ORFs (Samayoa, Yildiz, and Karplus 2011; Pauli, Valen, and Schier 2015). Exemplified by the identification of tens of sORFs (with supportive metadata), REPARATION’s utilization of Ribo-seq signal pattern at least in part alleviates the pitfalls of traditional bacterial gene prediction algorithms concerning the identification of sORFs.

Based on matching N-terminal proteomics evidence and the sequenced N-terminals from *Ecogene,* REPARATION accurately predicts 86 and 89% of the ORFs with experimental evidence. Overall, the high correlative second amino acid frequency patterns observed when comparing annotated versus re-annotated/new ORFs provide further proof of the accuracy and resolution of start codon selection in case of REPARATION predicted ORFs. Nonetheless, start site selection by REPARATION resulted in a loss of 6% of the N-terminal supported gene starts which exceeded the S-curve thresholds. While the existence of multiple N-terminal proteoforms in bacteria in contrast to the single ORF selection by REPARATION is likely the main explanatory reason for this inconsistency, the discrepancy between predicted and N-terminally supported start, might especially in the case of short truncations also be contributed (in part) by the lack of accuracy of start codon selection. REPARATION could potentially take advantage of improved measures or features to increase the prediction power of the classifier. At present REPARATION is the first attempt to perform a *de novo* putative ORF delineation in prokaryotic genomes that relies on Ribo-seq data. With automated bacterial gene prediction algorithms estimated to have false prediction rate of up to 30% (Angelova, Kalajdziski, and Kocarev 2010), machine learning algorithms that learn properties from Ribo-seq experiments such as REPARARTION pave the way for a more reliable (re-)annotation of prokaryotic genomes, a much desired need in the current era of (prokaryotic) genome sequencing.

## SOFTWARE

REPARATION software is available at https://github.com/Biobix/REPARATION.

## ACCESSION NUMBERS

Ribo-seq and RNA-seq sequencing data reported in this paper have been deposited in NCBI’s Gene Expression Omnibus with the accession number GSE91066.

All MS proteomics data and search results have been deposited to the ProteomeXchange Consortium via the PRIDE (Vizcaino et al. 2016) partner repository with the dataset identifier PXD005844 for the *Salmonella typhimurium SL1344* datasets and PXD005901 for the *E. coli* K12 str. MG1655 dataset. Reviewers can access the *Salmonella* datasets by using ‘’ as username and ‘Zg0VLXnS’ as password while the *E. coli* dataset can be accessed using ‘’ as username and ‘cn5cG4jW’ as password.

## ACKNOWLEDGMENTS

We thank Prof. Kris Gevaert for financial support of this research (Research Foundation - Flanders (FWO-Vlaanderen), project number G.0440.10). P.V.D. acknowledges support from the Research Foundation - Flanders (FWO-Vlaanderen), project number G.0269.13N. G.M. is a postdoctoral fellow at the Research Foundation – Flanders (FWO-Vlaanderen).

## AUTHOR CONTRIBUTIONS

E.N., A.G., E.V., G.M. and P.V.D. conceived the study; E.N., G.M. and P.V.D. wrote the manuscript; E.N. performed the computational analysis; P.V.D. performed the proteomics experiments, E.N. and P.V.D. performed the proteomics analyses; V.J. and P.V.D. prepared the Ribo-seq libraries. G.M. and P.V.D. supervised the research.

## COMPETING FINANCIAL INTERESTS

The authors declare that they have no competing financial interests.

